# Enhancer of Zeste Homolog 2 (*Ezh2*) is essential for patterning of multiple musculoskeletal tissues but dispensable for tendon differentiation

**DOI:** 10.1101/2020.10.30.361949

**Authors:** Deepanwita Pal, Scott M. Riester, Bashar Hasan, Sara F. Tufa, Amel Dudakovic, Douglas R. Keene, Andre J. van Wijnen, Ronen Schweitzer

## Abstract

An efficient musculoskeletal system depends on the precise assembly and coordinated growth and function of muscles, skeleton and tendons. However, the mechanisms that drive integrated musculoskeletal development and coordinated growth and differentiation of each of these tissues are still being uncovered. Epigenetic modifiers have emerged as critical regulators of cell fate differentiation, but so far almost nothing is known about their roles in tendon biology. Previous studies have shown that epigenetic modifications driven by Enhancer of zeste homolog 2 (EZH2), a major histone methyltransferase, have significant roles in vertebrate development including skeletal patterning and bone formation. We now find that targeting *Ezh2* through the limb mesenchyme also has significant effects on tendon and muscle patterning, likely reflecting the essential roles of early mesenchymal cues mediated by *Ezh2* for coordinated patterning and development of all tissues of the musculoskeletal system. Conversely, loss of *Ezh2* in the tendon cells did not disrupt the tendon cell fate suggesting that tenocyte differentiation and tendon maturation are independent of *Ezh2* signaling.

## Introduction

Epigenetic regulation comprises of heritable changes in gene expression that do not alter the underlying DNA sequence but instead rely on adapting the chromatin (1). Predominant epigenetic mechanisms include DNA methylation in CpG-rich promoters, non-coding RNAs and covalent modifications on specific residues in histones such as acetylation and methylation among others (2). Histone methylation is widely accepted as a key component of the epigenetic machinery that contributes to both stability and reversibility of gene expression (3) and has been associated with development and homeostasis (4). While histone methylation occurs on all basic residues, lysine methylations are the most extensively characterized (1). Methylation enzymes catalyze the addition of methyl groups donated from S-adenosylmethionine to histones and comprise of three families – (i) Su(var)3–9, Enhancer of Zeste, Trithorax (SET) – domain containing proteins, (ii) DOT1 (disruptor of telomeric silencing) - like proteins (iii) arginine N-methyltransferase (PRMT) proteins. Although originally thought to be permanent modifications, the discovery of lysine-specific demethylase 1A revealed the reversibility of histone methylation (5). The dynamic nature and reversibility of histone modifiers represents tremendous potential for therapeutic approaches and drug discovery, making them attractive targets for pharmacotherapies including orthopedic treatments (6, 7).

Breakthrough discoveries over the last decade have transformed our knowledge of epigenetic mechanisms in development and disease [8]. However, analysis of the function of epigenetic regulators in musculoskeletal biology is still evolving. In this work, we focused on a major histone methyltransferase called Enhancer of zeste homolog 2 (EZH2), a mammalian homolog of the Drosophila protein E(Z) (originally termed Enx-1) (8). *Ezh2* is the catalytic subunit of the Polycomb repressive complex (PRC2) and represents a highly evolutionarily conserved Polycomb group (PcG) member [10]. The functional relevance of the PcG proteins is highlighted by the numerous developmental defects reported in mice deficient for these proteins (9–11). *Ezh2* is distinct among the PcG proteins due to its evolutionarily conserved SET domain, which catalyzes histone H3 lysine (K) 27 methylation, eventually leading to transcriptional repression. *Ezh2* is important for early vertebrate development, as is evident from the phenotype of mice lacking *Ezh2*, which die around gastrulation (12). In zebrafish, *Ezh2* activity is dispensable for tissue specification but required for the maintenance of differentiated cell fates in the heart, liver and pancreas (13). *Ezh2* is also required for neural crest-derived bone and cartilage formation, with severe craniofacial defects observed upon *Ezh2* depletion in neural crest cells (14). In limb development, *Ezh2* regulates anteroposterior patterning via maintenance of *Hox* gene expression (15). Loss of *Ezh2* in uncommitted limb mesenchymal cells leads to skeletal malformations and defective bone formation, reflecting key roles for *Ezh2* in osteoblast maturation and skeletal development (16–18). Because of the epigenetic role of *Ezh2* in several principal stages of development, it is likely that it has essential roles in the development of other musculoskeletal tissues, but this hypothesis has not yet been experimentally validated.

The musculoskeletal system is a coordinated assembly of the skeleton, muscles and tendons, where the tendons transmit the force of muscle contraction to the skeleton thereby enabling joint movement. We recently established a model of musculoskeletal integration in the developing embryo (19) using expression of *Scleraxis* (Scx), a transcription factor, which serves as a distinctive marker for tendon cells from progenitor stages through development (20). Our work demonstrated the considerable interdependence of musculoskeletal tissues for coordinated growth and tissue formation. Considering the significant role of *Ezh2* in cell fate determination for various tissues including osteogenic differentiation (17, 18), we hypothesized that it may also play a role in cell fate determination of muscles and tendons. The goal of this study was to test if tendons are affected by the epigenetic signaling of *Ezh2* and whether *Ezh2* is involved in tenogenic differentiation.

We find that loss of *Ezh2* in limb mesenchyme resulted in dramatic changes to muscle and tendon pattern likely reflecting early mesenchymal signals required for patterning and coordinated growth of the musculoskeletal system. Even though *Ezh2* is required for tendon patterning, we find that the regulation of tenocyte differentiation and maturation does not depend on *Ezh2* signaling.

## Materials and methods

### Mice Breeding Scheme and Animal Welfare

Existing mouse lines used in these studies were described previously: *Ezh2*^f/f^ (21), *Prx1*^Cre^ (22), *ScxGFP* tendon reporter (23). Direct targeting of *Ezh2*, to determine tendon specific effects, was performed using the tendon specific deletor *Scx*^Cre^ (24). All animal procedures were approved by the Institutional Animal Care and Use Committee at the Oregon Health & Science University and the Mayo Clinic and are consistent with animal care guidelines.

### X-ray imaging

To measure differences in skeletal development, X-ray imaging was employed to assess gross anatomical structure at 3-4 weeks. Both males and females were analyzed to account for gender specific effects.

### Skeletal preparations

Cartilage and bones of the forelimbs harvested from mice were visualized after staining with Alcian blue (Sigma) and Alizarin red S (Sigma) and subsequent clarification of soft tissue with KOH (25).

### Tissue Harvest

Mice were euthanized by carbon dioxide asphyxiation followed by thoracotomy. Forelimbs were resected as follows: a small slit in the skin near the scapula was made. Then the skin was lifted using forceps and a slit was made through the length of the limb on the dorsal side. This superficial slit through the skin was made up to the metacarpal-phalangeal joint in the digits, to enable better fixation of the tissues inside.

### Histology

For whole-mount visualization, tissues were skinned and images were captured on an MZFLIII dissecting microscope (Leica) with an EOS50D camera (Canon). Patterning of limb tendons and muscles was acquired in serial transverse sections of 12 μm (26) and immunostaining was performed as previously described (27). Tendons were visualized using *ScxGFP* expression or anti-collagen I (ColI) antibody (Southern Biotech). Antigen retrieval using citrate buffer for the ColI staining was performed using the PELCO Biowave microwave (Ted Pella, Inc). A monoclonal antibody for My32 (Sigma Aldrich) was used to detect muscle-specific Type II myosin heavy chain (MHC). Images were captured using Zeiss ApoTome2 on AxioImager (Zeiss). Image processing was carried out using Zeiss Zen software.

### Transmission electron microscopy (TEM)

P21 and P28 mouse limbs were skinned and fixed in 1.5% glutaraldehyde/1.5% paraformaldehyde (Tousimis Research Corporation) in Dulbecco’s serum-free media (DMEM) containing 0.05% tannic acid at 4°C for 1-2 weeks. Forelimbs were then rinsed in DMEM followed by extensive decalcification in 0.2 M EDTA in 50mM Tris in a Pelco 3450 microwave (Ted Pella, Inc) at 94.5 W. Tissues were fixed again in 1.5% glutaraldehyde/1.5% paraformaldehyde with 0.05% tannic acid over night at 4°C, rinsed in DMEM, then post-fixated in 1% OsO4 overnight. Samples were washed in DMEM, dehydrated in a graded series of ethanol to 100%, rinsed in propylene oxide, and finally embedded in Spurrs epoxy. Tendons of interest were identified by collecting 1 μm sections stained with an epoxy tissue stain (Electron Microscopy Sciences). Ultrathin sections containing tendons of interest were cut at 80 nm, contrasted with uranyl acetate and lead citrate, and viewed with an FEI G20 TEM operated at 120 kV using an AMT 2 x 2K camera. TEM images of transverse sections were collected at several magnifications to enable morphological visualization of the collagen fibrils and gross tendon appearance.

## Results

### Deletion of *Ezh2* in limb mesenchyme affects tendon and musculoskeletal patterning

It was previously reported that global knockout of *Ezh2* in mice results in embryonic lethality (12) precluding analysis of specific roles in tendon development. Hence, for initial analysis, we targeted *Ezh2* in limb mesenchyme using *Prx1*^Cre^ (22). Previous studies established a role for *Ezh2* in anterior posterior patterning of the limb (15) and associated loss of *Ezh2* with several skeletal abnormalities including decreased vertebral height, shortened limbs and prematurely fused cranial structures (15, 16, 28). However, the effects of the loss of *Ezh2* on patterning and differentiation of the soft musculoskeletal tissues were not examined in these reports. Consistent with previous studies, we found that *Ezh2^Prx1Cre^* mutant limbs were considerably shorter compared to WT littermates and displayed anteroposterior patterning defects (15) (Fig 1). *Ezh2^Prx1Cre^* mutants also displayed movement limitations likely reflecting broad disruptions to the musculoskeletal system in addition to the reported skeletal defects (data not shown).

**Fig 1.**
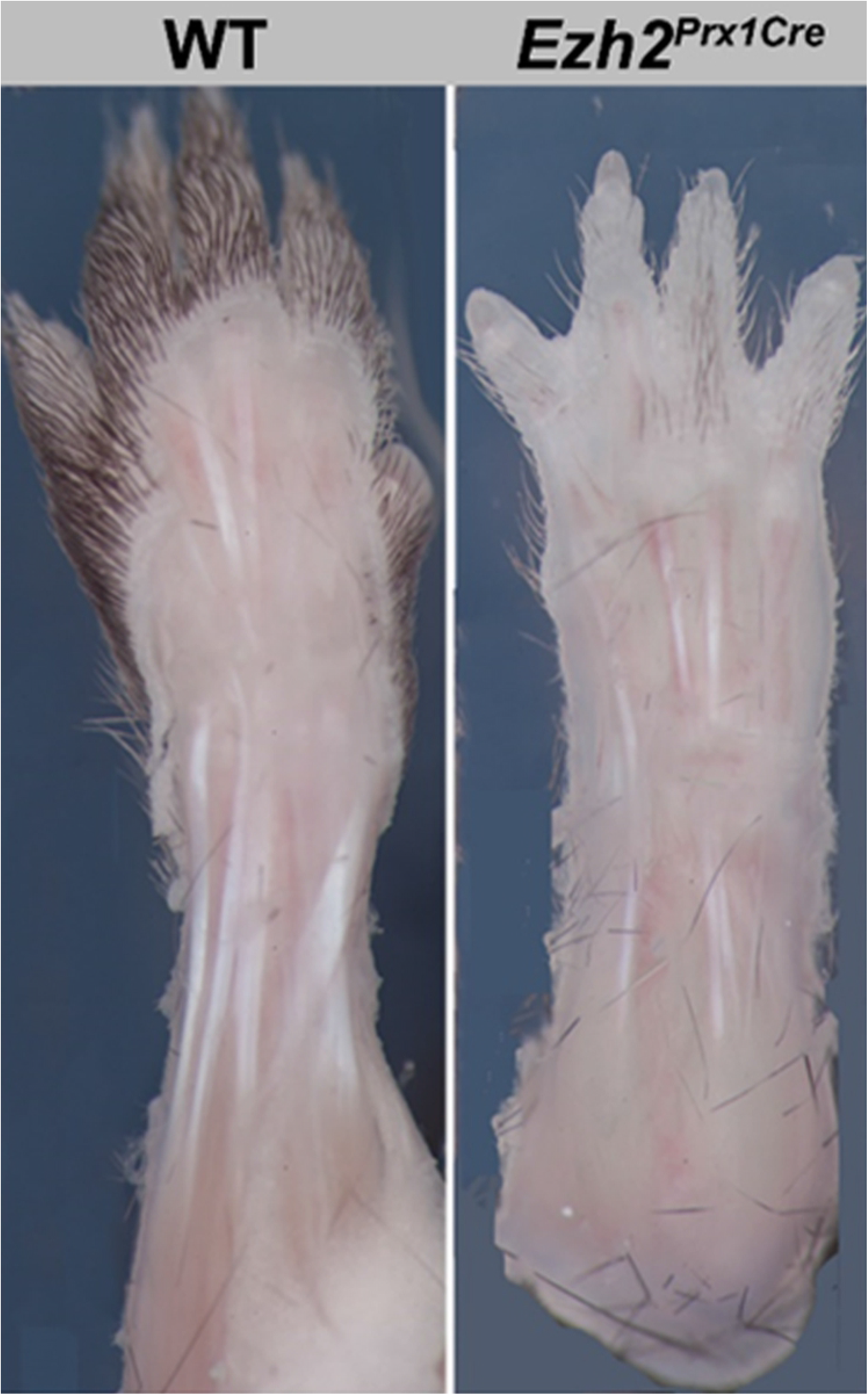
Deletion of Ezh2 in the limb mesenchyme affects skeletal patterning. Whole mount images of WT and *Ezh2^Prx1Cre^* limbs at 3 weeks

To examine the effects of targeting *Ezh2* in limb mesenchyme on the soft musculoskeletal tissues, the tendons and muscles, we stained for Collagen I (ColI; encoded by *Col1a1* or *Col1a2*) and Myosin Heavy Chain (MHC; encoded by *Myh1*) respectively (Fig 2B). The patterning of both the tendons and muscles was substantially affected in mutant limbs. Paw movement is controlled by two broad categories of muscles, intrinsic muscles whose muscle bellies are found within the paw and extrinsic muscles with muscle bellies that reside in the arm and long tendons that connect these muscles to their specific insertion points in the paw (26, 29).

**Fig 2.**
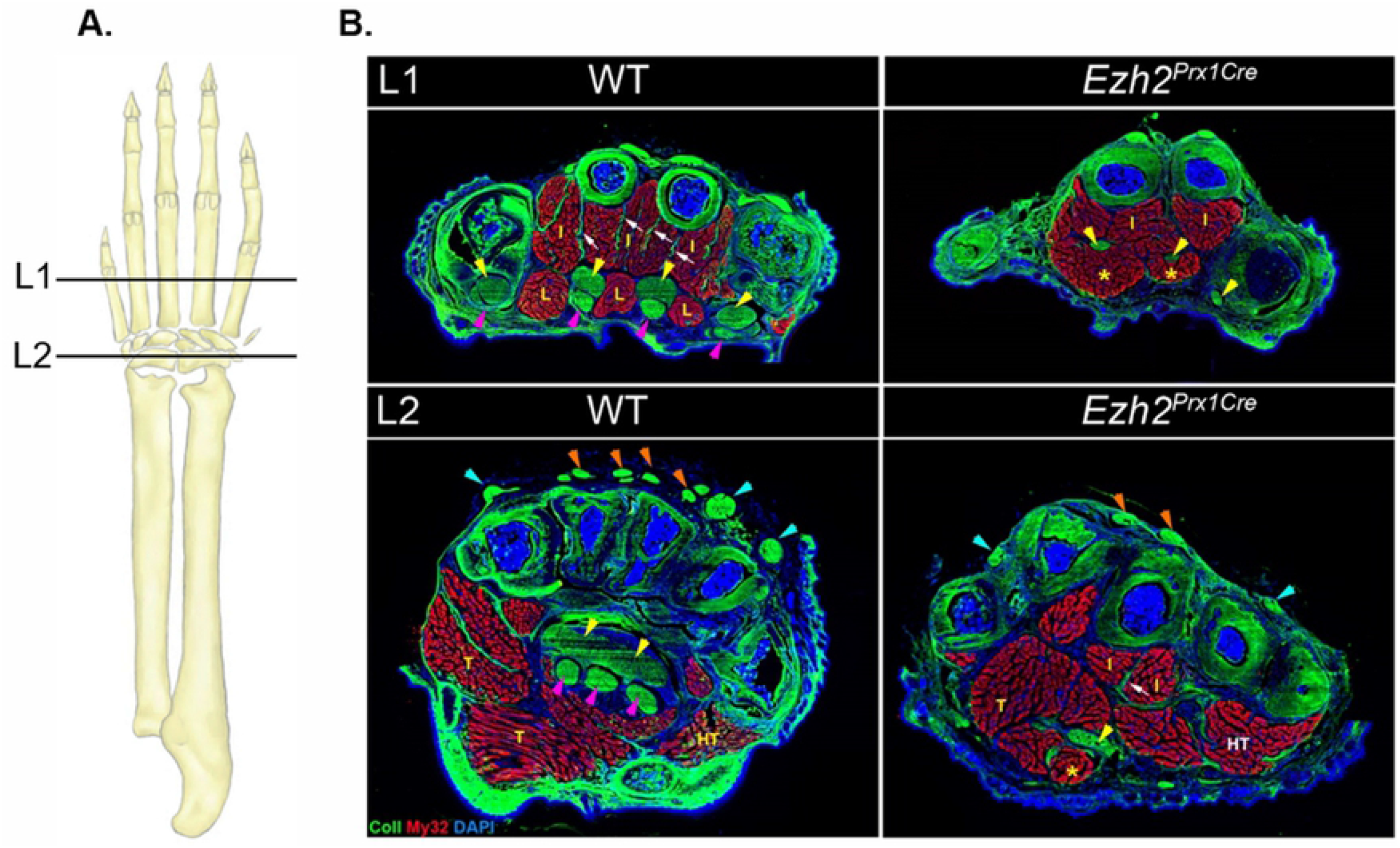
Deletion of Ezh2 in the limb mesenchyme affects muscle and tendon patterning. **A.** Schematic of the ventral side of the forelimb that illustrates the positions of the limb sections on the panels on the right. **B.** Images of transverse sections at the metacarpals (L1) and the wrist (L2) of WT and *Ezh2^Prx1Cre^* mutant limbs at P21. Coll staining was used to visualize tendons in WT and mutant limbs respectively while My32 staining was used for muscle. Yellow, pink, orange and teal arrowheads highlight the flexor digitorium profundus, flexor digitorium superficialis, extensor digitorium communis and the lateral extensor tendons respectively. White arrows indicate interosseous tendons. I, L, T and HT indicate the interosseous, lumbricals, thenar and hypothenar muscles respectively.

All extensor muscles are extrinsic and their tendons were severely disrupted in mutant forelimbs. Extensor communis tendons extend through the length of the digits and the skeletal insertions for other extensors are found in various carpal or metacarpal bones (26). Only two communis extensor tendons were detected in mutant limbs (Fig 2B, orange arrowheads) and most of the other lateral extensor tendons were missing in the mutant (Fig 2B, teal arrowheads). The major flexor muscles are the flexor digitorium profundus (FDP) extrinsic muscles. The FDP tendons are fused in the wrist and carpal regions and separate to individual tendons that insert in the distal part of each digit (26, 29). These FDP tendons, the most robust paw tendons, were significantly smaller throughout the paw of mutant forelimbs (Fig 2B, yellow arrowheads). The two major intrinsic groups of paw muscles are the interosseous muscles located directly ventral to the metacarpal bones and the lumbrical muscles that extend through the length of the metacarpals directly between FDP tendons ((26) and Fig 2B). The interosseous muscles were also significantly smaller and mis-patterned in mutant limbs (Fig 2B, I). Moreover, interosseous tendons could not be detected in distal sections and only a small subset of these tendons were detected more proximally (Fig 2B, white arrows). Mutant effects were even more dramatic on the lumbrical muscles. Lumbrical muscles have a distinct morphology between the FDP tendons in sections at metacarpal level ((26) and Fig 2B) and these muscles were completely absent in mutant paws (Fig 2B, L).

The severe effects of the loss of *Ezh2* on musculoskeletal patterning were further reflected in the fate of the flexor digitorium superficialis (FDS) muscles and tendons. While FDS muscles are extrinsic to the paw, they follow a unique developmental program. The FDS muscles first differentiate with multinucleated myofibers in the forepaw and subsequently translocate out of the paw and into the arm leaving behind the FDS tendons at their distal ends (30). The translocation of FDS muscles involves dramatic muscle elongation at the proximal end with concurrent retraction from the distal end and both of these processes did not occur in the *Ezh2^Prx1Cre^* mutant. In mutant sections, we found muscles directly underneath the FDP tendons instead of the FDS tendons that usually occupy this space (Fig 2B, yellow asterisks). We previously identified such a configuration in the limbs of *Scx* and paralyzed mutants in which FDS muscle translocation failed (27).

Together, these results demonstrate that the loss of *Ezh2* had a profound effect on the patterning, organization and size of muscles and tendons. Combined with previous reports of significant disruptions of skeletal patterning in *Ezh2^Prx1Cre^* mutants, these results suggest an essential role for *Ezh2* in overall musculoskeletal patterning.

### *Ezh2* is dispensable for the tendon cell fate and tenocyte maturation

*Ezh2* was implicated in cell fate differentiation and maturation and in addition to the early effects on skeletal development, *Ezh2* has essential roles during osteogenesis (17). Since tendons were greatly affected in the *Ezh2^Prx1Cre^* mutant, we investigated whether Ezh2 is also involved in tenocyte differentiation and the regulation of tendon growth and maturation. To examine *Ezh2* function directly in the tenocyte lineage, we used a tendon specific deletor *Scx^Cre^* (24) to target *Ezh2* specifically in tenocytes. First, we wanted to determine if skeletal patterning or growth was affected in these mutants, to exclude the possibility of secondary effects of skeletal disruptions on the tendons. Contrary to the *Ezh2^Prx1Cre^* mutant, skeletal pattern was not affected in *Ezh2^ScxCre^* mutants as evident from whole mount images, skeletal preparations and X-ray images (Fig 3). Furthermore, using the *ScxGFP* tendon reporter (23), we found that in mutant limbs the tendon system was largely intact and there were no significant differences between four week old WT and *Ezh2^ScxCre^* mutant littermates in tendon patterning (Fig 4B and C).

**Fig 3.**
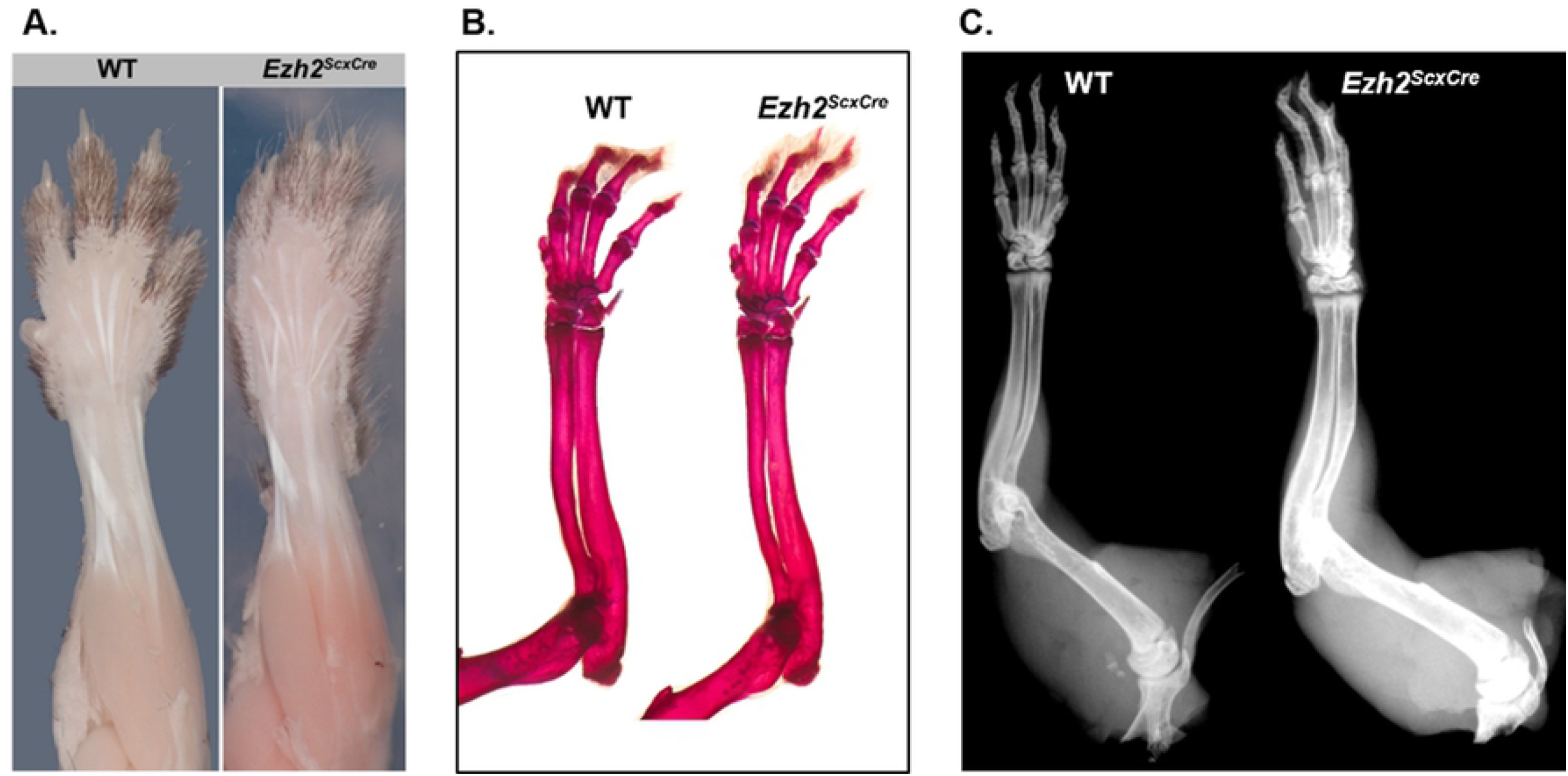
Targeted deletion of Ezh2 in tendons did not result in defects in skeletal patterning. Whole mount images **(A)** and skeletal preparations **(B)** and X-ray images **(C)** of WT and *Ezh2^ScxCre^* limbs at 4 weeks.

**Fig 4.**
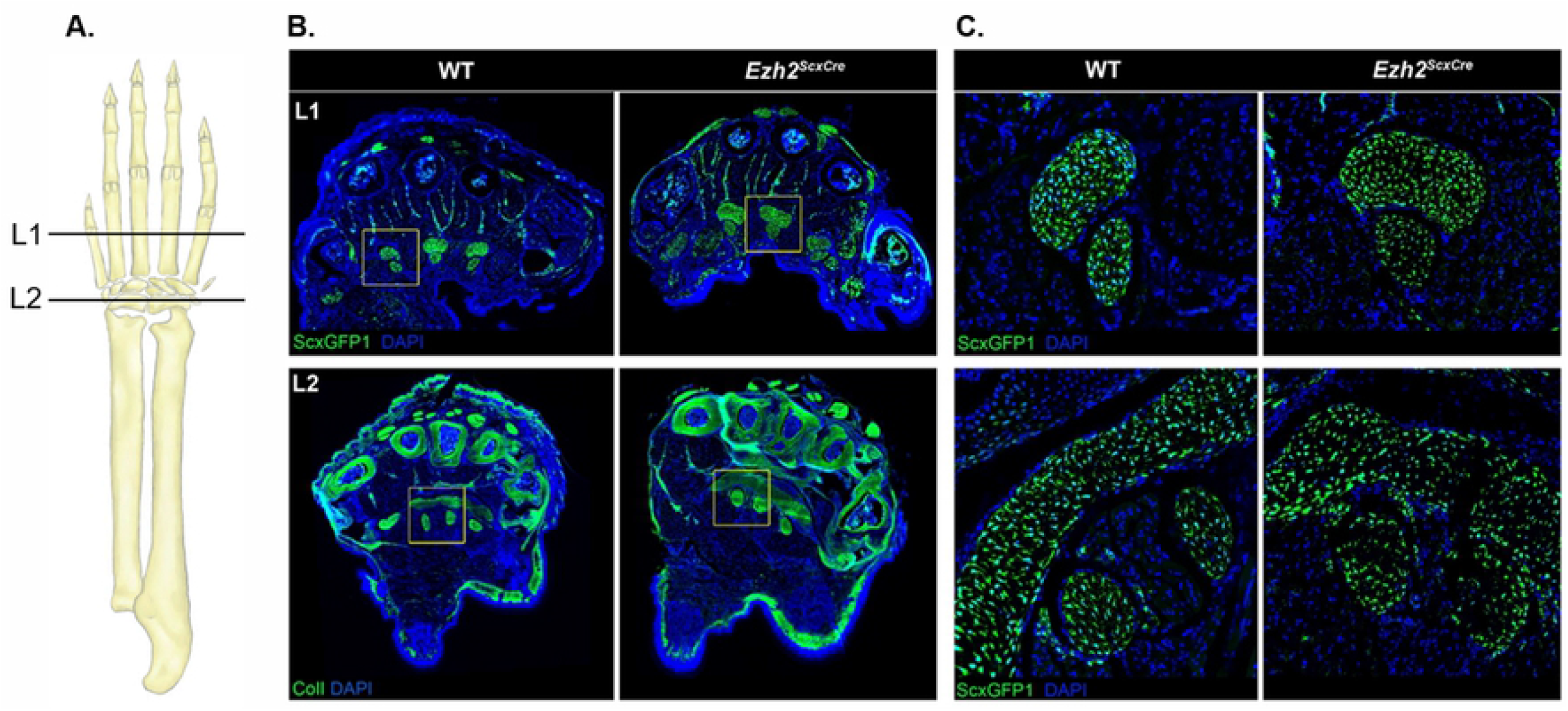
Loss of Ezh2 in tendon lineage cells did not affect tendon patterning. **A.** Schematic of the ventral side of the forelimb that illustrates the positions of the limb sections on the panels on the right. **B.** Images of transverse sections of WT and *Ezh2^ScxCre^* mutant at 4 weeks showing ScxGFP and ColI expression at the metacarpals (L1) and the wrist (L2). **C.** High magnification of the regions defined by the insets on the left showing transverse sections of the flexors of the WT and *Ezh2^ScxCre^* mutant.

Although we did not observe any tendon patterning defects in the *Scx^Cre^* mutant, we next examined the involvement of *Ezh2* in tendon differentiation. The characteristic feature of a mature tendon is the highly organized and robust collagen matrix. The collagen matrix occupies most of the tendon volume and it is composed of an assembly of collagen fibrils with heterogeneous cross section area that extend in parallel arrays through the length of the tendon (31). The pattern and quantitative parameters of the collagen matrix are commonly examined in high magnification transmission electron microscope (TEM) images. To determine if the tendon cell fate was affected in *Ezh2^ScxCre^* mutants, we therefore analyzed the organization of the collagen matrix in these mutants. We find that the collagen matrix was not disrupted in the tendons of *Ezh2^ScxCre^* mutants and fibril organization, shape and the distribution of fibril diameters all appear similar in comparison of TEM images from section of mutant and WT littermate tendons (Fig 5A).

**Fig 5.**
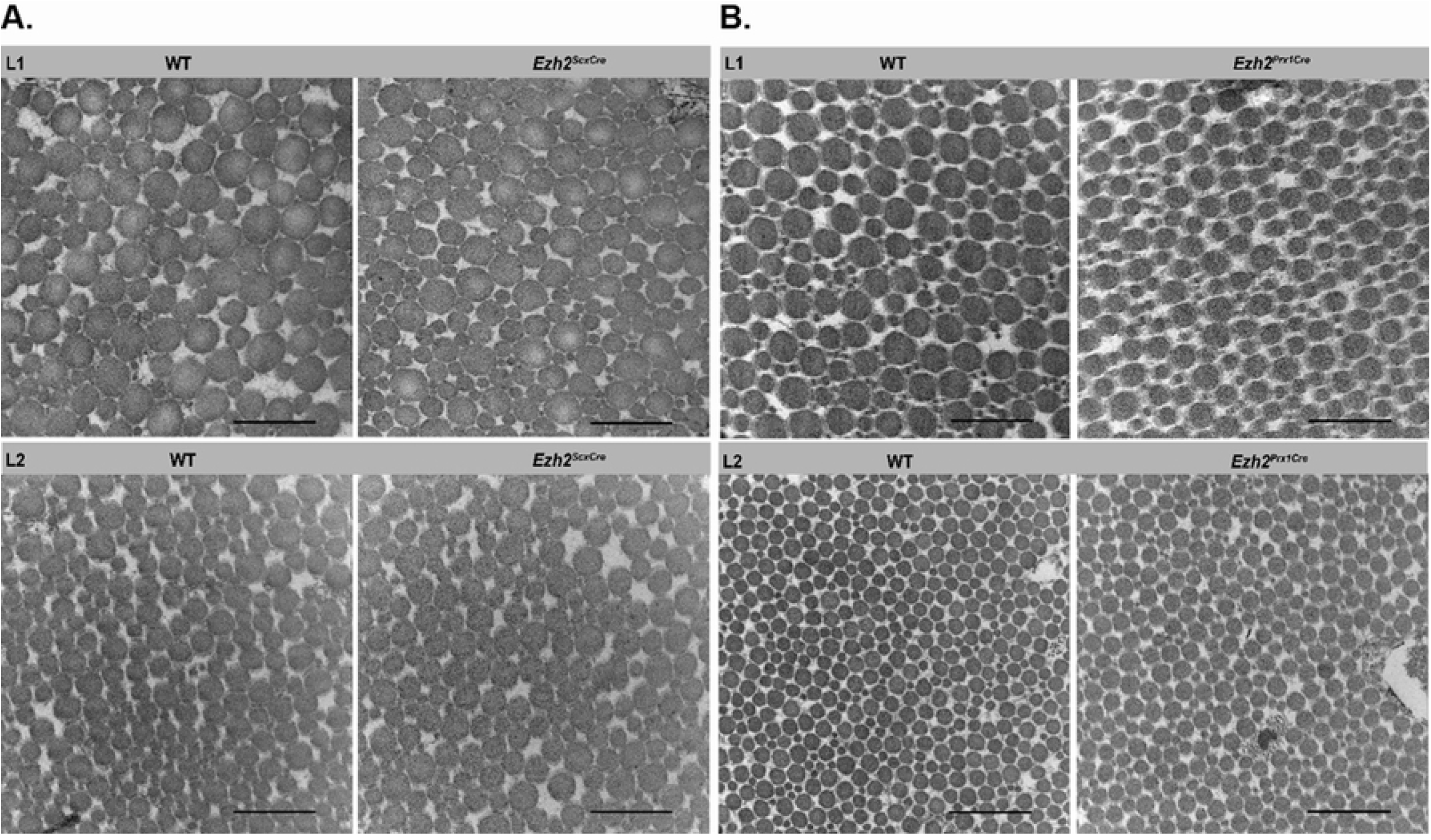
Ezh2 is dispensable for tendon cell fate and tenocyte maturation. **A.** TEM images of WT and *Ezh2^ScxCre^* mutant showing collagen structure in transverse sections of flexor digitorium profundus tendon in the digits (L1) and extensor carpi ulnaris in the wrist (L2). Scale: 500nm, Magnification: 50,000X. **B.** TEM images of WT and *Ezh2^Prx1Cre^* mutant showing collagen structure in transverse sections through flexor digitorium profundus tendon in the digits (L1) and extensor carpi radialis brevis tendon in the wrist (L2). Scale: 500nm, Magnification: 50,000X.

The onset of *Prx1^Cre^* activity in early limb mesenchyme at E9.5 precedes recombination by the *Scx^Cre^* driver that emerges in tendon cells around E13.5. Since the tendons and all other tissues of the musculoskeletal system were profoundly affected in *Ezh2^Prx1Cre^* mutants, we investigated whether early targeting of EZH2 in the *Ezh2^Prx1Cre^* mutants had a more significant effect on the tendon cell fate. We therefore also analyzed the collagen matrix in tendons from these mutants. However, we found that despite the considerable disruption of tendon pattern and size in the *Ezh2^Prx1Cre^* mutants, the loss of *Ezh2* had no effect on tendon maturation and differentiation and the collagen matrix in mutant tendon was similar to that of WT littermates (Fig 5B). Taken together, our results suggest that while *Ezh2* activity is essential in early stages of limb development for tendon and overall musculoskeletal patterning, *Ezh2* activity is dispensable within the tenocyte-lineage and thus not essential for the regulation of the tendon cell fate.

## Discussion

In this study, we have examined the role of the histone methyltransferase *Ezh2* in tendon development. Our findings show that *Ezh2*-mediated signals are important for early tendon and overall musculoskeletal patterning, but *Ezh2* is not essential for the regulation of tendon cell fate.

It was previously reported that *Ezh2* plays an essential role in early limb patterning (15). The role of *Ezh2* in very early aspects of limb patterning was further highlighted by the fact that anterior-posterior patterning was more dramatically affected when *Ezh2* was targeted with *T^Cre^* before the initiation of the limb bud compared with the outcome of targeting *Ezh2* with *Prx1^Cre^* that becomes active only at the initial stages of limb bud formation (15). In agreement with this, we consistently found in *Ezh2^Prx1Cre^*, substantial disruption of skeletal patterning and loss of digit 1 ((16) and Fig 1).

The developmental regulatory signals that underlie the specific pattern of the cartilage condensations in the developing embryo remain obscure and even less is known about the initial cues that guide muscle or tendon patterning. However, skeletal patterning is independent of the presence of muscles and the basic layout of muscle pattern can be detected even in a skeletal-less limb (19, 24). Early patterning of all the musculoskeletal tissues may therefore be guided by signals from the limb mesenchyme (32, 33). It is not yet known what these signals are or if all musculoskeletal tissues are affected by the same signals or if there are distinct cues for each of these tissues. It was therefore interesting to examine the effects of the loss of *Ezh2* on muscle and tendon patterning. Remarkably, the tendons and the muscles were also severely affected in the *Ezh2^Prx1Cre^* mutant, suggesting that *Ezh2* activity underlies basic aspects of mesenchymal patterning that guide the patterning and development of all musculoskeletal tissues.

The loss of *Ezh2* in limb mesenchyme resulted in severe and complex effects on tendon development. Various extensor and flexor tendons were missing in mutant limbs and the existing tendons were drastically smaller that their WT counterparts, an effect seen most dramatically in the FDP tendons (Fig 2B). Contrary to the independent development of cartilage and muscles, tendon development is inherently dependent on the other musculoskeletal tissues (19, 24, 34, 35). We and others have previously demonstrated that tendon induction and development within the autopod is dependent on cartilage and tendon development in the zeugopod is dependent on the presence of muscles (19, 35). Because of the interdependence of tendon development in the paw on signals from the cartilage, the failure of tendon development may reflect an indirect effect through disruption of signal(s) that emanate from the cartilage to regulate tendon formation. On the other hand, the observed tendon defects may also reflect a direct role for *Ezh2* in tendon formation by regulating specific mesenchymal cues that guide or instruct tendon development. It is not possible at this time to determine if *Ezh2* activity plays a direct or indirect role in the development of these tendons.

The complex role of *Ezh2* activity in regulation of muscle and tendon development is especially reflected in the effects on flexor muscles and tendon. FDP tendons were significantly smaller, FDS tendons were missing in the mutant paw and FDS muscles were found in the paw instead of their normal position in the arm. We previously described a unique developmental path for the FDS muscles and tendons (30). The FDS muscles first organize as fully formed muscles in the paw. They subsequently elongate proximally and translocate from the paw and into the arm. The FDS tendons are formed in the wake of the FDS muscles as they recede from the paw (30). In mutants that disrupt tendon development (*Scx*) or muscle activity (*mdg*) muscle translocation is disrupted and the FDS muscles remain within the paw, resulting also in failure of FDS tendon development (19, 27). The FDS muscle and tendon phenotype in *Ezh2^Prx1Cre^* mutant limbs may thus reflect a direct failure of muscle activity or tendon development. It may however also reflect the absence of other cues that initiate, support or propel FDS muscle movement.

Notably, failures in tendon growth could also reflect a direct role for *Ezh2* in the regulation of tendon growth. However, the tendon phenotypes observed in *Ezh2^Prx1Cre^* mutant limbs were not recapitulated when *Ezh2* was targeted with the tendon-specific deletor, *Scx^Cre^*, suggesting that *Ezh2* is most likely essential for early patterning of all musculoskeletal tissues. However, we cannot rule out the possibility that *Ezh2* has an early role in limb mesenchyme, which is later essential for tendon differentiation and growth. Since the onset of *Scx^Cre^* activity is later than that of *Prx1^Cre^*, such an early function may be bypassed in the *Ezh2^ScxCre^* mutant. Finally, it is well established that tendon growth is dependent on biomechanical activity (19). Considering the skeletal patterning disruptions in *Ezh2^Prx1Cre^* mutants and the mobility issues they experience, it is conceivable that some aspects of the tendon phenotypes may be attributed to restricted biomechanical activity.

*Ezh2* activity has been implicated in cell fate determination and differentiation (36), as well as maintenance of the stem cell state in the mesenchymal linages (37).The complexity of these functions of *Ezh2* was uncovered in studies of the skeletal tissues. Loss of *Ezh2* in the osteogenic lineage resulted in reduced bone formation, but conditional loss of *Ezh2* in uncommitted mesenchymal cells yields skeletal patterning defects, including shortened forelimbs, craniosynostosis and clinodactyly (17). Conversely, loss of *Ezh2* in chondrocytes did not disrupt cartilage development despite the appearance of osteogenic gene expression in the mutant chondrocytes (38). We therefore examined if *Ezh2* has a role in tenogenic differentiation, but did not find any effect on the normal production of the prototypic extracellular matrix of tendons in mutant tendons (Fig 5).

Finally, in analysis of the consequences of targeting the *Ezh2* gene it is important to consider the possible roles of functional homologs that may substitute to provide the same molecular activity in different tissues or at different stages of development and maturation. Indeed, *Ezh1*, a functional homolog of *Ezh2*, is ubiquitously expressed whereas *Ezh2* expression is associated mostly with proliferating tissues (39). Hence, it is possible that the homologs share redundant features that would allow *Ezh1* to compensate for the functions of *Ezh2* in various tissues including tendons. A recent report (40) indeed suggests that *Ezh1* and Ezh2 could potentially compensate for each other in skeletal development. It remains to be seen if such a mechanism is also active during tendon development.

## Acknowledgments

We would like to thank all past and present members of our laboratories, including Brian Pryce, Fuhua Xu, Emily Camilleri and Roman Thaler for stimulating discussions, providing reagents and generously sharing ideas.

## Author contributions

**Conceptualization**: Deepanwita Pal, Scott Riester, Amel Dudakovic, Andre van Wijnen, Ronen Schweitzer

**Data curation**: Deepanwita Pal

**Formal analysis**: Sara Tufa, Douglas Keene

**Funding acquisition**: Andre van Wijnen, Ronen Schweitzer

**Investigation**: Deepanwita Pal, Scott Riester, Bashar Hasan, Amel Dudakovic, Sara Tufa

**Methodology**: Deepanwita Pal, Scott Riester, Bashar Hasan, Amel Dudakovic, Sara Tufa, Douglas Keene, Andre van Wijnen, Ronen Schweitzer

**Project administration**: Andre van Wijnen, Ronen Schweitzer

**Resources**: Douglas Keene, Andre van Wijnen, Ronen Schweitzer

**Supervision**: Andre van Wijnen, Ronen Schweitzer

**Validation**: Deepanwita Pal, Scott Riester, Bashar Hasan, Amel Dudakovic, Andre van Wijnen, Ronen Schweitzer

**Visualization**: Deepanwita Pal, Amel Dudakovic, Andre van Wijnen, Ronen Schweitzer

**Writing – Original draft**: Deepanwita Pal, Ronen Schweitzer

**Writing – Review and editing**: Deepanwita Pal, Scott Riester, Bashar Hasan, Sara Tufa, Amel Dudakovic, Douglas Keene, Andre van Wijnen, Ronen Schweitzer

## References

1. Allis CD, Jenuwein T. The molecular hallmarks of epigenetic control. Nat Rev Genet. 2016;17(8):487–500.

2. Barrero MJ, Boue S, Izpisua Belmonte JC. Epigenetic mechanisms that regulate cell identity. Cell Stem Cell. 2010;7(5):565–70.

3. Martin C, Zhang Y. The diverse functions of histone lysine methylation. Nat Rev Mol Cell Biol. 2005;6(11):838–49.

4. Oppermann U. Why is epigenetics important in understanding the pathogenesis of inflammatory musculoskeletal diseases? Arthritis Res Ther. 2013;15(2):209.

5. Byvoet P, Shepherd GR, Hardin JM, Noland BJ. The distribution and turnover of labeled methyl groups in histone fractions of cultured mammalian cells. Arch Biochem Biophys. 1972;148(2):558–67.

6. Arrowsmith CH, Bountra C, Fish PV, Lee K, Schapira M. Epigenetic protein families: a new frontier for drug discovery. Nat Rev Drug Discov. 2012;11(5):384–400.

7. van Wijnen AJ, Westendorf JJ. Epigenetics as a New Frontier in Orthopedic Regenerative Medicine and Oncology. J Orthop Res. 2019;37(7):1465–74.

8. Hobert O, Jallal B, Ullrich A. Interaction of Vav with ENX-1, a putative transcriptional regulator of homeobox gene expression. Mol Cell Biol. 1996;16(6):3066–73.

9. Gould A. Functions of mammalian Polycomb group and trithorax group related genes. Curr Opin Genet Dev. 1997;7(4):488–94.

10. van der Lugt NM, Domen J, Linders K, van Roon M, Robanus-Maandag E, te Riele H, et al. Posterior transformation, neurological abnormalities, and severe hematopoietic defects in mice with a targeted deletion of the bmi-1 proto-oncogene. Genes Dev. 1994;8(7):757–69.

11. Dudakovic A, van Wijnen AJ. Epigenetic Control of Osteoblast Differentiation by Enhancer of Zeste Homolog 2 (EZH2). Current Molecular Biology Reports. 2017;3(2):94–106.

12. O’Carroll D, Erhardt S, Pagani M, Barton SC, Surani MA, Jenuwein T. The polycomb-group gene Ezh2 is required for early mouse development. Mol Cell Biol. 2001;21(13):4330–6.

13. San B, Chrispijn ND, Wittkopp N, van Heeringen SJ, Lagendijk AK, Aben M, et al. Normal formation of a vertebrate body plan and loss of tissue maintenance in the absence of ezh2. Sci Rep. 2016;6:24658.

14. Schwarz D, Varum S, Zemke M, Scholer A, Baggiolini A, Draganova K, et al. Ezh2 is required for neural crest-derived cartilage and bone formation. Development. 2014;141(4):867–77.

15. Wyngaarden LA, Delgado-Olguin P, Su IH, Bruneau BG, Hopyan S. Ezh2 regulates anteroposterior axis specification and proximodistal axis elongation in the developing limb. Development. 2011;138(17):3759–67.

16. Dudakovic A, Camilleri ET, Xu F, Riester SM, McGee-Lawrence ME, Bradley EW, et al. Epigenetic Control of Skeletal Development by the Histone Methyltransferase Ezh2. J Biol Chem. 2015;290(46):27604–17.

17. Dudakovic A, Camilleri ET, Paradise CR, Samsonraj RM, Gluscevic M, Paggi CA, et al. Enhancer of zeste homolog 2 (Ezh2) controls bone formation and cell cycle progression during osteogenesis in mice. J Biol Chem. 2018;293(33):12894–907.

18. Dudakovic A, Samsonraj RM, Paradise CR, Galeano-Garces C, Mol MO, Galeano-Garces D, et al. Inhibition of the epigenetic suppressor EZH2 primes osteogenic differentiation mediated by BMP2. J Biol Chem. 2020;295(23):7877–93.

19. Huang AH, Riordan TJ, Pryce B, Weibel JL, Watson SS, Long F, et al. Musculoskeletal integration at the wrist underlies the modular development of limb tendons. Development. 2015;142(14):2431–41.

20. Schweitzer R, Chyung JH, Murtaugh LC, Brent AE, Rosen V, Olson EN, et al. Analysis of the tendon cell fate using Scleraxis, a specific marker for tendons and ligaments. Development. 2001;128(19):3855–66.

21. Su IH, Basavaraj A, Krutchinsky AN, Hobert O, Ullrich A, Chait BT, et al. Ezh2 controls B cell development through histone H3 methylation and Igh rearrangement. Nat Immunol. 2003;4(2):124–31.

22. Logan M, Martin JF, Nagy A, Lobe C, Olson EN, Tabin CJ. Expression of Cre Recombinase in the developing mouse limb bud driven by a Prxl enhancer. Genesis. 2002;33(2):77–80.

23. Pryce BA, Brent AE, Murchison ND, Tabin CJ, Schweitzer R. Generation of transgenic tendon reporters, ScxGFP and ScxAP, using regulatory elements of the scleraxis gene. Dev Dyn. 2007;236(6):1677–82.

24. Blitz E, Viukov S, Sharir A, Shwartz Y, Galloway JL, Pryce BA, et al. Bone ridge patterning during musculoskeletal assembly is mediated through SCX regulation of Bmp4 at the tendon-skeleton junction. Dev Cell. 2009;17(6):861–73.

25. McLeod MJ. Differential staining of cartilage and bone in whole mouse fetuses by alcian blue and alizarin red S. Teratology. 1980;22(3):299–301.

26. Watson SS, Riordan TJ, Pryce BA, Schweitzer R. Tendons and muscles of the mouse forelimb during embryonic development. Dev Dyn. 2009;238(3):693–700.

27. Murchison ND, Price BA, Conner DA, Keene DR, Olson EN, Tabin CJ, et al. Regulation of tendon differentiation by scleraxis distinguishes force-transmitting tendons from muscle-anchoring tendons. Development. 2007;134(14):2697–708.

28. Hemming S, Cakouros D, Codrington J, Vandyke K, Arthur A, Zannettino A, et al. EZH2 deletion in early mesenchyme compromises postnatal bone microarchitecture and structural integrity and accelerates remodeling. FASEB J. 2017;31(3):1011–27.

29. Delaurier A, Burton N, Bennett M, Baldock R, Davidson D, Mohun TJ, et al. The Mouse Limb Anatomy Atlas: an interactive 3D tool for studying embryonic limb patterning. BMC Dev Biol. 2008;8:83.

30. Huang AH, Riordan TJ, Wang L, Eyal S, Zelzer E, Brigande JV, et al. Repositioning forelimb superficialis muscles: tendon attachment and muscle activity enable active relocation of functional myofibers. Dev Cell. 2013;26(5):544–51.

31. Benjamin M, Ralphs JR. Tendons and ligaments--an overview. Histol Histopathol. 1997;12(4):1135–44.

32. Hasson P, DeLaurier A, Bennett M, Grigorieva E, Naiche LA, Papaioannou VE, et al. Tbx4 and tbx5 acting in connective tissue are required for limb muscle and tendon patterning. Dev Cell. 2010;18(1):148–56.

33. Colasanto MP, Eyal S, Mohassel P, Bamshad M, Bonnemann CG, Zelzer E, et al. Development of a subset of forelimb muscles and their attachment sites requires the ulnar-mammary syndrome gene Tbx3. Dis Model Mech. 2016;9(11):1257–69.

34. Edom-Vovard F, Duprez D. Signals regulating tendon formation during chick embryonic development. Dev Dyn. 2004;229(3):449–57.

35. Kardon G. Muscle and tendon morphogenesis in the avian hind limb. Development. 1998;125(20):4019–32.

36. Chou RH, Yu YL, Hung MC. The roles of EZH2 in cell lineage commitment. Am J Transl Res. 2011;3(3):243–50.

37. Sen B, Paradise CR, Xie Z, Sankaran J, Uzer G, Styner M, et al. beta-Catenin Preserves the Stem State of Murine Bone Marrow Stromal Cells Through Activation of EZH2. J Bone Miner Res. 2020;35(6):1149–62.

38. Camilleri ET, Dudakovic A, Riester SM, Galeano-Garces C, Paradise CR, Bradley EW, et al. Loss of histone methyltransferase Ezh2 stimulates an osteogenic transcriptional program in chondrocytes but does not affect cartilage development. J Biol Chem. 2018;293(49):19001–11.

39. Margueron R, Li G, Sarma K, Blais A, Zavadil J, Woodcock CL, et al. Ezh1 and Ezh2 maintain repressive chromatin through different mechanisms. Mol Cell. 2008;32(4):503–18.

40. Lui JC, Garrison P, Nguyen Q, Ad M, Keembiyehetty C, Chen W, et al. EZH1 and EZH2 promote skeletal growth by repressing inhibitors of chondrocyte proliferation and hypertrophy. Nat Commun. 2016;7:13685.

